# Novel rabies virus variant for bi-directional optical control reveals modulatory influence of the pulvinar on visual cortex in rat

**DOI:** 10.1101/2020.07.23.218610

**Authors:** LR Scholl, L Zhang, AT Foik, DC Lyon

## Abstract

Optogenetic tools have become of great utility in the causal analysis of systems in the brain. However, current optogenetic techniques do not reliably support both excitation and suppression of the same cells in vivo, limiting analysis and slowing research. Here we developed a novel glycoprotein-deleted rabies virus expressing two channelrhodopsin proteins, GtACR2 and Chrimson, in order to independently manipulate excitatory and inhibitory transmembrane potentials, respectively. Using this approach, we demonstrated that rodent pulvinar neurons modulate cortical size tuning and suppress flash responses, but do not drive activity in visual cortex. While our goal was primarily to develop this novel method to study the structure-function organization of thalamocortical circuits, this technique is readily applicable to study any brain region.

---

Manipulating neural activity is a highly effective way to deduce the role of complex cortical networks on sensory perception and behavior. More crude methods of ablation and lesion have given way to small molecule receptor agonists and electrical microstimulation, and, more recently, to optogenetic proteins that are reversible, do not interfere with the normal physiology of cells, and can quickly and efficiently raise or lower membrane potentials (Deisseroth, 2011; Yizhar et al. 2011; Bernstein et al. 2012; Prigge et al. 2012). These recent optogenetic methods can reach high spatial and temporal resolutions making it possible to dissect cortical network connections and related functions on a fine scale (e.g., Adesnik et al. 2012; Lee et al. 2012; Olsen et al. 2012; Wilson et al. 2012; Xue et al. 2014). Even so, light-sensitive proteins only allow either anions or cations to flow across the neuronal membrane, inducing either depolarization or hyperpolarization, but not both. Introducing a single activating or inhibiting opsin does not provide absolute control over cell activity, making it impossible to drive or suppress downstream activity in all situations due to complex excitatory and inhibitory network structures. In addition, continuous excitation or inhibition can lead to adaptation and reduced efficacy (Lignani et al., 2013), and is impractical over the long-term because the balance between excitation and inhibition that is important for network dynamics (van Vreeswijk & Sompolinsky, 1996) is altered.

One system where these issues are apparent is the projections between the thalamus, in particular the pulvinar, and visual cortex. The pulvinar nucleus, called the lateral posterior nucleus (LP) in rodents, is known to have influence over activity in visual cortex (Saalmann et al., 2012; Tohmi et al., 2014; Zhou et al., 2016), but not much is known about what information is being transmitted (Roth et al., 2015; Zhou et al., 2017). The pulvinar receives diverse input (see Figure 1a), ranging from attention signals from frontal cortex (Romanski et al., 1997; Wilke et al., 2009) to input from multiple sensory cortical areas (Kaas & Lyon, 2007; Kamishina et al., 2009; Juavinett et al., 2019) and superior colliculus (Benevento & Standage, 1983; Takahashi, 1985; Nakamura et al., 2015), to projections from melanopsin-containing retinal ganglion cells (Allen et al., 2016). It is unlikely that a nucleus with such diverse inputs has only one function, or that neurons in the pulvinar all have similar responses to the same stimulus. This makes it difficult to know in advance whether activation or inactivation (or both) of pulvinar neurons will have an effect on cortical systems. Furthermore, projections from the thalamus can target both excitatory and inhibitory subnetworks (Bruno & Sakmann, 2006; Cruikshank et al., 2007), so although thalamocortical projections are typically excitatory in nature (Reid & Alonso, 1995; Gil et al., 1999), the same population of pulvinar cells might have a facilitating drive over one cortical system but an inhibitory drive over another.

**Figure 1.**
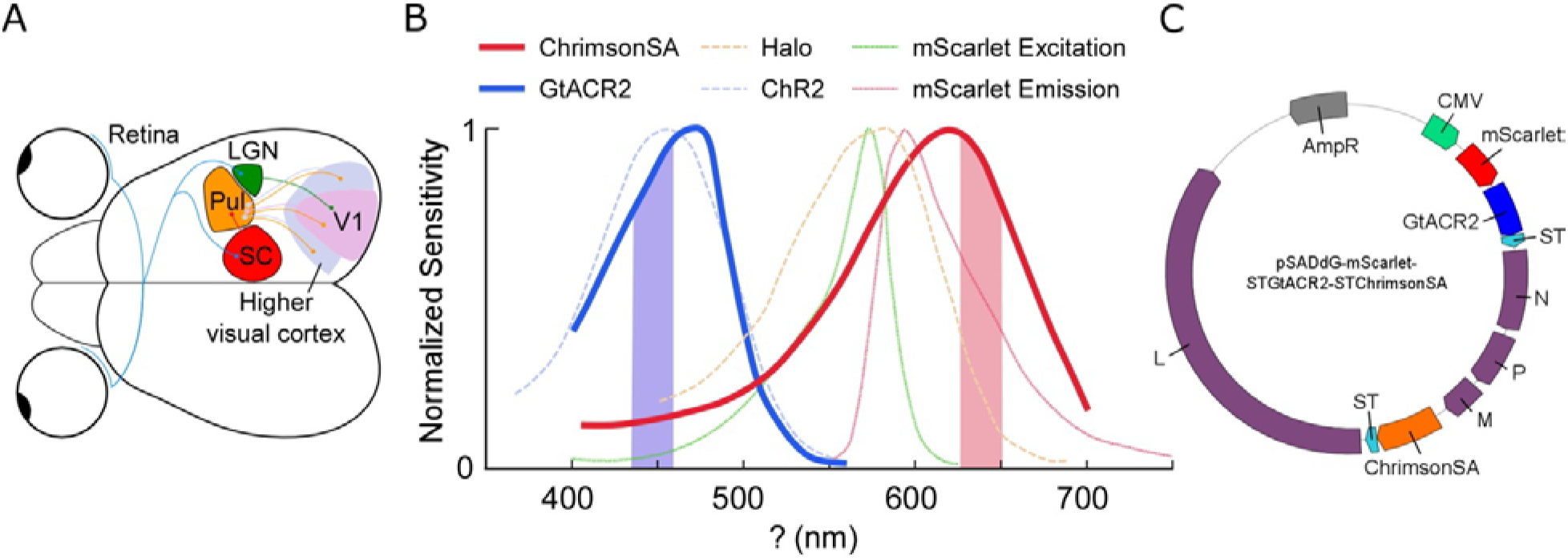
Rodent visual system overview and virus design. (a) Retinal ganglion cells project to the lateral geniculate nucleus (LGN) and the superior colliculus (SC). Most of the pulvinar (Pul) receives driving input from cortex, including primary (V1) and higher visual cortex, but the caudal part also receives input from SC. Projections from the pulvinar mainly target higher visual cortex, with more sparse connections to V1. Not to scale.(b) Excitation spectra for Chrimson (solid orange; adapted from Klapoetke et al., 2014) and GtACR2 (solid blue; adapted from Govorunova et al., 2015). Halorhodopsin (Halo; dotted orange) and Channelrhodopsin-2 (ChR2; dotted blue) are included for reference (Adapted from Han & Boyden, 2007), as are the excitation and emission spectra for the fluorescent protein mScarlet (dotted green and red lines; adapted from Bindels et al., 2016). The overlap between GtACR2 and Chrimson excitation spectra is similar to the Halo-ChR2 overlap, which Han & Boyden (2007) have shown to work independently in the same cell. (c) Plasmid encoding the modified rabies virus genome. mScarlet and GtACR2 transgenes were cloned immediately following the transcription start sequence with additional M/G intergenic region sequences between each coding region. ChrimsonSA was cloned in place of G. Soma-targeting sequences (ST) were fused to GtACR2 and ChrimsonSA to prevent axonal activation.

Here we introduce a method to retrogradely deliver a pair of independently activated light-gated ion channels, GtACR2 and Chrimson, with selectivity for positively and negatively charged ions, respectively. With this new tool, we probe the inputs to visual cortex from the pulvinar to ask what modulatory effect pulvinar has on visually evoked responses in cortex. Since we can perform both activation and inactivation of projection neurons, we effectively cut experiment times in half, and increase statistical power by studying a single population. We also demonstrate the effectiveness of the virus by recording local cortical activity in response to multi wavelength laser activation.

## Materials and Methods

### Bi-Directional Optical Construct Design

In order to independently excite and inhibit neurons, we selected two channelrhodopsin proteins with compatible spectral and electrochemical characteristics. These are Chrimson, a fast red-shifted channelrhodopsin variant (Klapoetke et al., 2014; Oda et al., 2018) and GtACR2, a blue-shifted mutant from a class of anion-channelrhodopsins (Govorunova et al., 2015). It has been demonstrated previously that Chrimson effectively induces cation flow and elicits neural spiking activity, while GtACR2 selectively passes anions and suppresses neural spiking. The combined absorption spectra allows a yellow colored laser around 630 nm to exclusively activate Chrimson but not GtACR2, and a blue colored laser around 450 nm to activate GtACR2, with only residual activation of Chrimson (Figure 1b). Critically, this residual activation is overcome by the higher sensitivity of GtACR2 compared to Chrimson, such that with suitable laser power, GtACR2-driven anion flow should dominate. This design is superior to any utilizing light-driven anion pumps, such as halorhodopsin (Halo), due to the millisecond timescale and high sensitivity of both channelrhodopsins, and offers greater spectral separation than other approaches, such as between ChR2 and Halo (see Figure 1b; Han & Boyden, 2007; Zhang et al., 2007).

In order to visualize traced neurons *in vivo* or in histological slides, a fluorescent protein compatible with the two channelrhodopsins’ absorption spectrums is needed. We employed the fluorescent protein mScarlet (Bindels et al. 2016); this reporter is suitable not only because of its near-infrared emission spectrum, giving the added benefit of having deep tissue penetration for in vivo imaging, but also because it is by far the brightest red fluorescent protein reported (Bindels et al., 2016). Moreover, the excitation spectrum of mScarlet avoids both laser wavelengths selected to activate Chimson and GtACR2, preventing any unwanted fluorescence during optogenetic manipulation (Figure 1b).

To package the effector genes we used glycoprotein-deleted rabies virus (RVdG), which offers several advantages over other methods of gene delivery (Figure 1c; Wickersham et al. 2007a; Osakada et al. 2011). When enveloped with rabies glycoprotein or optimized glycoprotein (oG; Kim et al. 2016), RVdG is able to retrogradely trace and deliver genes to the cell body of all pre-synaptically connected neurons, revealing the projections of one brain area to another (e.g., Connolly et al. 2012; Negwer et al. 2017; Foik et al. 2018). In addition, a major benefit of using rabies virus to package channelrhodopsin genes is the ability to pseudotype the virus (Wickersham et al. 2007b) to target specific protein receptors delivered transgenically, via helper viruses, and through single cell electroporation (Marshel et al. 2010; Wall et al. 2010; Miyamichi et al. 2011; Rancz et al. 2011; Kim et al. 2015; Callaway and Luo, 2015; Wertz et al. 2015; Wall et al., 2016). For example, recently we developed a suite of helper viruses designed to transduce avian tumor virus receptor A (TVA) and oG to excitatory (LV-αCamKII) or inhibitory (AAV-GAD1) subpopulations (Liu et al., 2013; Lean et al., 2019). As such, RVdG pseudotyped with the ASLV-A envelope glycoprotein (EnvA) will selectively infect neurons expressing the TVA receptor, and the oG delivered in trans will enable monosynaptic infection of retrogradely connected neurons.

Several more considerations were made in designing the viral genome. Two measures were taken to reduce the likelihood of interactions between the opins. First, a faster variant of the Chrimson protein, ChrimsonSA (Oda et al., 2018), was chosen due to its further red-shifted spectra and reduced sensitivity to light, thus increasing the relative sensitivity of the anion-channelrhodopsin. To further increase GtACR2 dominance in the short-wavelength range, the transgenes were arranged such that GtACR2 was located near the beginning of the genome, leading to increased transcription efficiency (see Figure 1c; Finke et al., 2000; Schnell et al., 2010). Finally, in order to restrict the channelrhodopsins to the cell soma, preventing unwanted axonal activation, both channelrhodopsins were fused with the soma-targeting signal (ST) from the voltage-gated potassium channel Kv2.1 (Lim et al., 2000; Mahn et al., 2018).

### Virus production

Virus was prepared following the protocol by Osakada and Callaway (2013) and pseudotyped with oG following Ciabatti et al. (2017). Plasmid synthesis, sequence verification, and maxiprep were performed by Gene Universal Inc (Newark, DE). BHK cells expressing rabies glycoprotein SADB19G (B7GG, provided by the Callaway laboratory) were transfected with the genomic plasmid pSADdG-mScarlet-STGtACR2-STChrimsonSA in addition to plasmids encoding rabies viral proteins N, L, P, and G using Lipofectamine 2000 (Invitrogen, Waltham, MA) transfection reagent. Following transfection, the 100 mm dish was incubated for 6 d at 3% CO2 and 35°C, then transferred to a 150 mm dish and incubated for a further 6 d before the supernatant was removed and passed through a 0.45 μm polyethersulfone filter.

BHK cells expressing optimized rabies glycoprotein (TGoG, provided by the Tripodi laboratory) were infected with 5 ml of viral supernatant and maintained at 3% CO2 and 35°C for 1 d, then washed thoroughly and transferred to a clean dish for another 5-6 d to produce viral supernatant with only oG coated virus. This supernatant was subsequently used to infect five 150 mm dishes of the same cell line in order to amplify the virus. Supernatant was collected twice during incubation at 3% CO2 and 35°C after 6 and 10 days, then filtered and transferred to an ultracentrifuge (rotor SW28, Beckman Coulter) for 2 h at 19,400 g and 4°C. Purified virus was resuspended in phosphate buffered saline for 1 h at 4°C before 2% fetal bovine serum was added. Aliquots for injection were stored at −80°C. Titer was assessed by infecting HEK 293T cells (Sigma-Aldrich) with serial dilutions of modified virus, to ensure at least ~1 × 10^9^ infectious units/ mL was achieved. Figure 2a shows infected 293T cells during titration.

**Figure 2.**
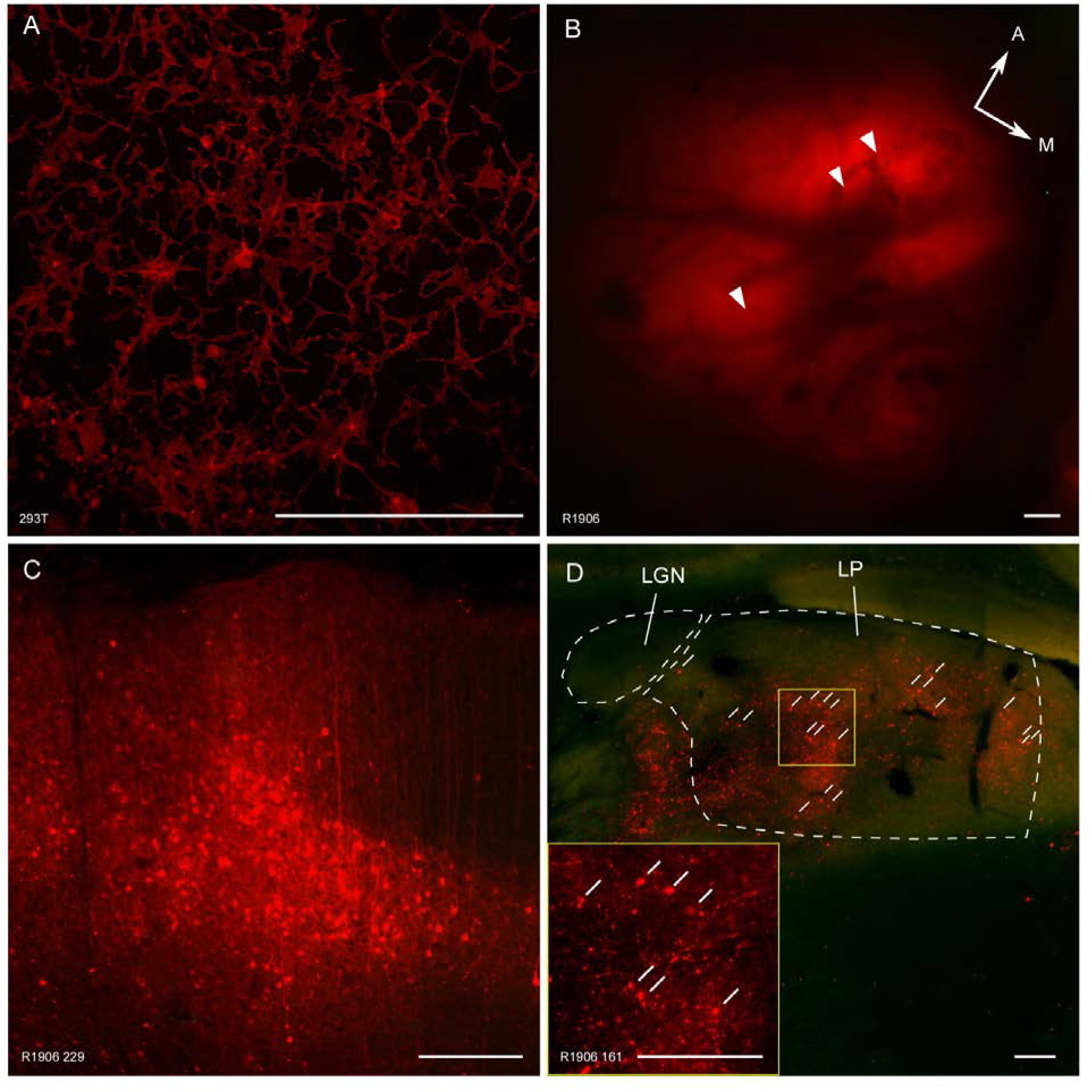
mScarlet fluorescence in cultured cells and rat neurons. (a) Infected 293T cells, (b) *in vivo* fluorescence one week after cortical injections (arrowheads), and the same case *ex vivo* with labeled neurons in cortex (c) and in LP (d). Scale bars equal 250 μm

### Surgery and *Electrophysiology*

Injections of modified rabies virus were carried out in four adult female Long-Evans rats. All procedures were approved by the University of California, Irvine Institutional Animal Care and Use Committee and the Institutional Biosafety Committee, and followed the guidelines of the National Institutes of Health. Prior to surgery, rats were initially anesthetized with 2% isoflurane in a mixture of 30% oxygen and 70% nitrous oxide, and maintained with 1 to 1.5% isoflurane in the same mixture. Using a stereotaxic apparatus, a craniotomy was performed to expose the caudal neocortex of one hemisphere. A glass micropipette was cut to approximately 20 μm diameter, filled with rabies virus suspension, and lowered into the brain using a motorized microdrive. Multiple injections (spaced ~1 mm) of rabies virus were made in medial or lateral V2 at several depths ranging from 200 to 1600 μm. Following a 6-7 d survival period to allow for infection and expression of transgenes, an eight-channel linear probe (U-probe; Plexon, Dallas, TX) was lowered into the injected site guided by in vivo fluorescent imaging (see Figure 2c).

Multichannel recordings were acquired using a 32-channel Scout recording system (Ripple, UT, USA). Local field potentials (LFP) were captured at 1 kHz sampling frequency from signals filtered between 0.3 to 250 Hz and with 60 Hz noise removed. Signals containing spikes were bandpass filtered from 500 Hz to 7 kHz and stored at 30 kHz sampling frequency. Spikes were sorted online in Trellis software (Ripple, UT, USA) while performing visual stimulation. For LFP recordings, only electrodes more than 250 um apart were considered for analysis (Katzner et al., 2009).

### Optical and Visual Stimuli

Laser stimulation was delivered via a 100 μm diameter optic fiber positioned at the surface of the injection site for cortical activation or at the apex of the lateral posterior nucleus for thalamic activation (stereotaxic coordinates −4 mm from bregma, 1.75 to 2.75 mm lateral). Two diode lasers housed in a beam combiner (Omicron, Dudenhofen, Germany) coupled to the optic fiber, one 594 nm yellow laser (Mambo; Cobolt, Stockholm, Sweden) and one 473 nm blue laser (LuxX 473; Omicron) were triggered using the parallel port and synchronized within 10 ms of visual stimuli onset. To excite neurons with Chrimson, the 594 nm laser was turned on at 20 mW and either 5, 10, or 20 Hz frequency (square wave, 50% duty cycle) for 1 second. GtACR2 was targeted to inhibit cellular activity using the 473 nm laser at 5 mW and 10 Hz frequency for 1 second.

Visual stimuli were generated in Experica (http://experica.org) and displayed on a gamma-corrected LCD monitor (55 inches, 60 Hz; RCA, New York, NY) at 1920×1080 pixels resolution and 52 cd/m^2^ mean luminance. Stimulus onset times were corrected for LCD monitor response time using a photodiode and microcontroller. Flashes of white and black filling the display were used to test pulvinar’s influence on V2 activity. Transitions from black to white were pseudo-randomly paired with either 594 or 473 nm wavelength laser light, or with no laser light. Visually responsive cells in V2 were found using 100% contrast drifting sinusoidal gratings. Cells were first tested for optimal orientation, spatial frequency, temporal frequency, contrast, and receptive field location. Next, using optimal parameters cells were tested with stimulus aperture sizes ranging from 0 to 80º of visual angle while psuedo-randomly paired with 594 or 473 nm wavelength laser light, or no laser light.

### Data analysis and statistics

Visually evoked potentials (VEP) were averaged across 50 trials for laser and flash stimuli. Peri-stimulus time histograms (PSTH) were constructed using 20 ms bin widths and averaged across 50 trials for laser and flash stimuli, or 20 trials for grating stimuli. Mean firing rate (MFR) was calculated for spikes in the first 200 ms following stimulus presentation. Average data are presented as mean ± standard error of the mean (SEM) unless otherwise noted. For cells tested with grating stimuli, optimal size was defined as the diameter with the maximum MFR under the no laser condition. Suppressed size was the diameter bigger than the optimal size with the lowest MFR. Wilcoxon signed rank tests were used to test the effects of each laser to baseline and between the two laser wavelengths. Pearson’s correlations were used to compare electrode depth to the difference in mean VEP or MFR between 594 nm and 473 nm wavelength laser conditions. Values of P ≤ 0.05 were considered significant for all tests.

### Histology

Following several recording sessions, but no more than 12 d post injection, rats were deeply anesthetized with Euthasol and transcardially perfused first with saline, then with 4% paraformaldehyde in phosphate buffered saline. Brains were removed and cryoprotected in 30% sucrose for at least 24 hours, then sectioned coronally on a freezing microtome to 40 μm thickness and mounted on glass microscope slides in polyvinyl alcohol mounting medium with 1,4-diazabicyclo-octane (PVA-DABCO, prepared in-house).

Sections were scanned using a fluorescent microscope (Axioplan 2; Zeiss, White Plains, NY) equipped with a 10x objective and motorized stage. Images were captured with a monochromatic low-noise CCD camera (Sensicam qe, PCO AG, Kelheim, Germany) and corrected for lamp misalignment by dividing each pixel by corresponding pixels in a flat field image acquired for each color channel. Corrected images were stitched using stage coordinates with regions of 10 overlapping pixels between images in which average pixel values were used. Neurons were identified based on the presence of the cell soma and dendrites. Fluorescently labeled neurons were then annotated by anatomical brain region based on the rat brain atlas by Paxinos and Watson (2013). Image correction and stitching were performed in MATLAB using the multisection-imager toolbox (http://github.com/leoscholl/multisection-imager).

## Results

The genomic plasmid pSADdG-mScarlet-STGtACR2-STChrimsonSA was successfully synthesized and its sequence verified before virus production. Following transfection, the virus was pseudotyped with optimized glycoprotein (oG) in order to increase infection efficiency (Kim et al. 2016) and titer was verified in the 293T cell line (Figure 2a). The virus was injected into V2, revealing bright *in vivo* (Figure 2b) and *ex vivo* (Figure 2c) mScarlet fluorescence. Retrogradely labeled cells were visible not only locally near the injection site (Figure 2c) but also in the thalamus (Figure 2d), in particular the lateral posterior nucleus (LP).

Extracellular cortical recordings in V2 were made *in vivo* during simultaneous local laser manipulation (Figure 3). Single units recorded near the injection site were excited by 5 Hz yellow laser stimulation at the brain surface and suppressed by 5 Hz blue laser stimulation (Figure 4), and were reliably excitable at and above 10 Hz laser stimulation (square wave, 50% duty cycle; Figures 4 and 5). For subsequent tests, a frequency of 10 Hz was used for both laser wavelengths.

**Figure 3.**
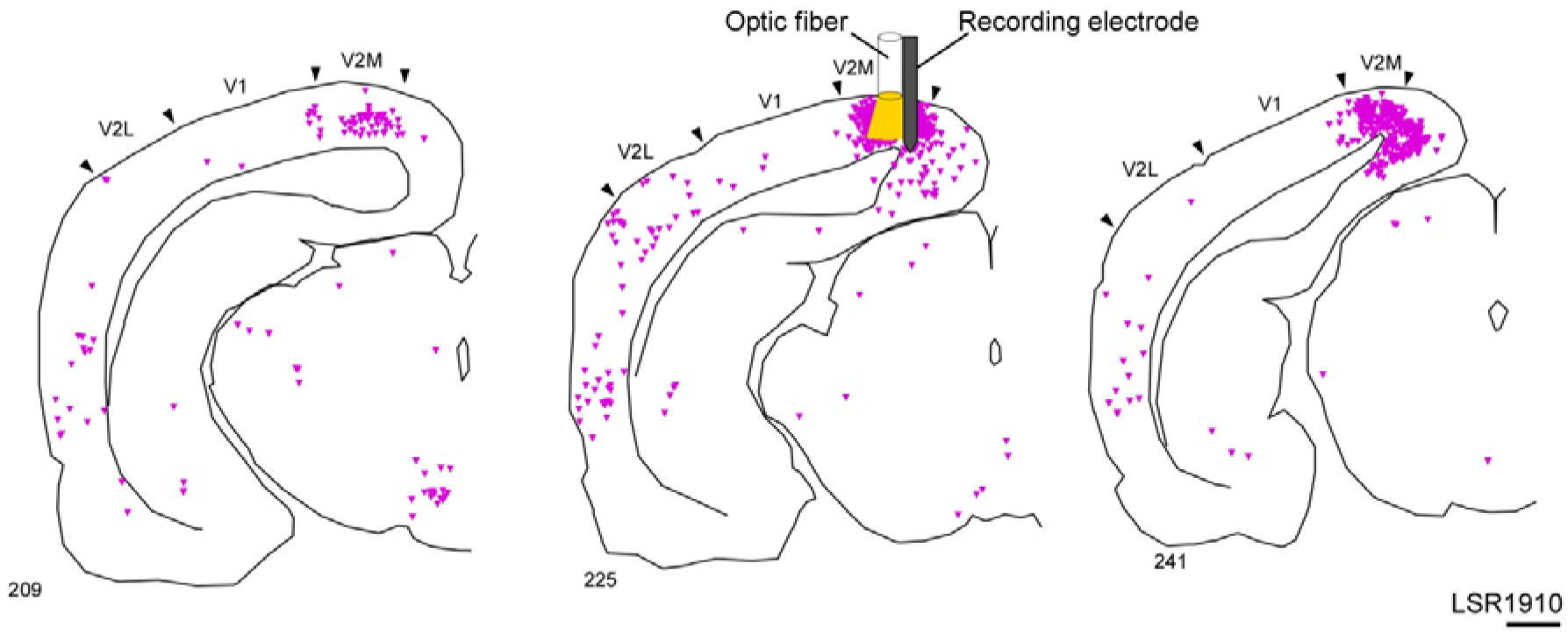
Local V2 neurons targeted for optical manipulation. Reconstruction of case LSR1910 in which mScarlet-STGtACR2-STChrimsonSA rabies virus was injected into medial visual cortex where subsequent recordings and laser activation were performed. Scale bar equals 1 mm.

**Figure 4.**
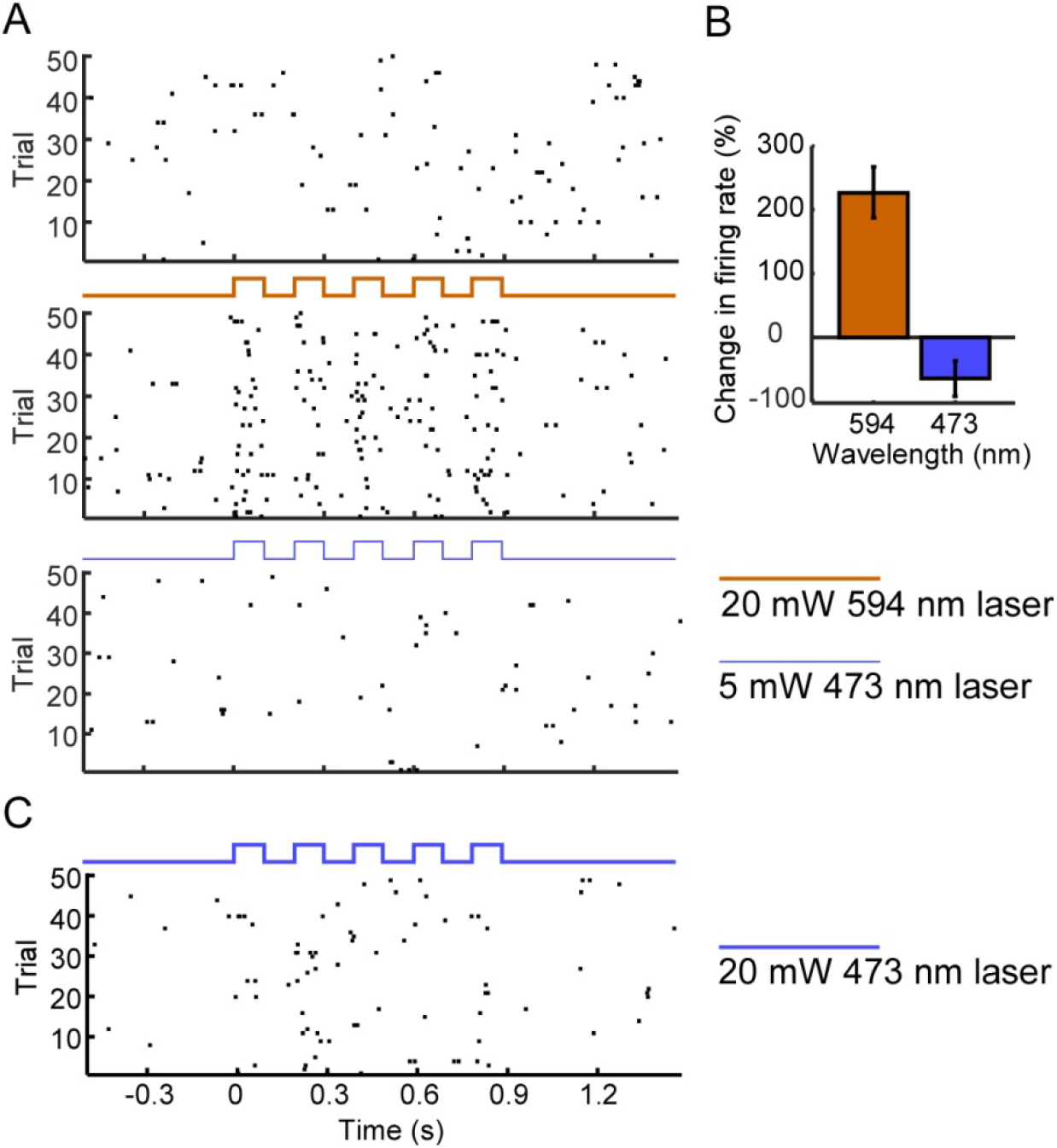
Cortical excitation with 594 nm and inhibition with 473 nm wavelength lasers with no visual stimulus. The firing rate of this cell is elevated over 200% under yellow laser light at 20 mW, and decreased by 75% under blue laser light at 5mW. (a) Raster plots of each laser condition. Black marks indicate spikes during trials with no laser stimulation, yellow marks indicate spikes during yellow laser stimulation at 5 Hz, and blue marks indicate spikes during blue laser stimulation at 5 Hz. (b) Summary of the percent change in firing rate from baseline (no laser) to yellow (594 nm) or blue (473 nm) laser excitation. (c) Example of the same cell being excited by 20 mW blue laser light.

**Figure 5.**
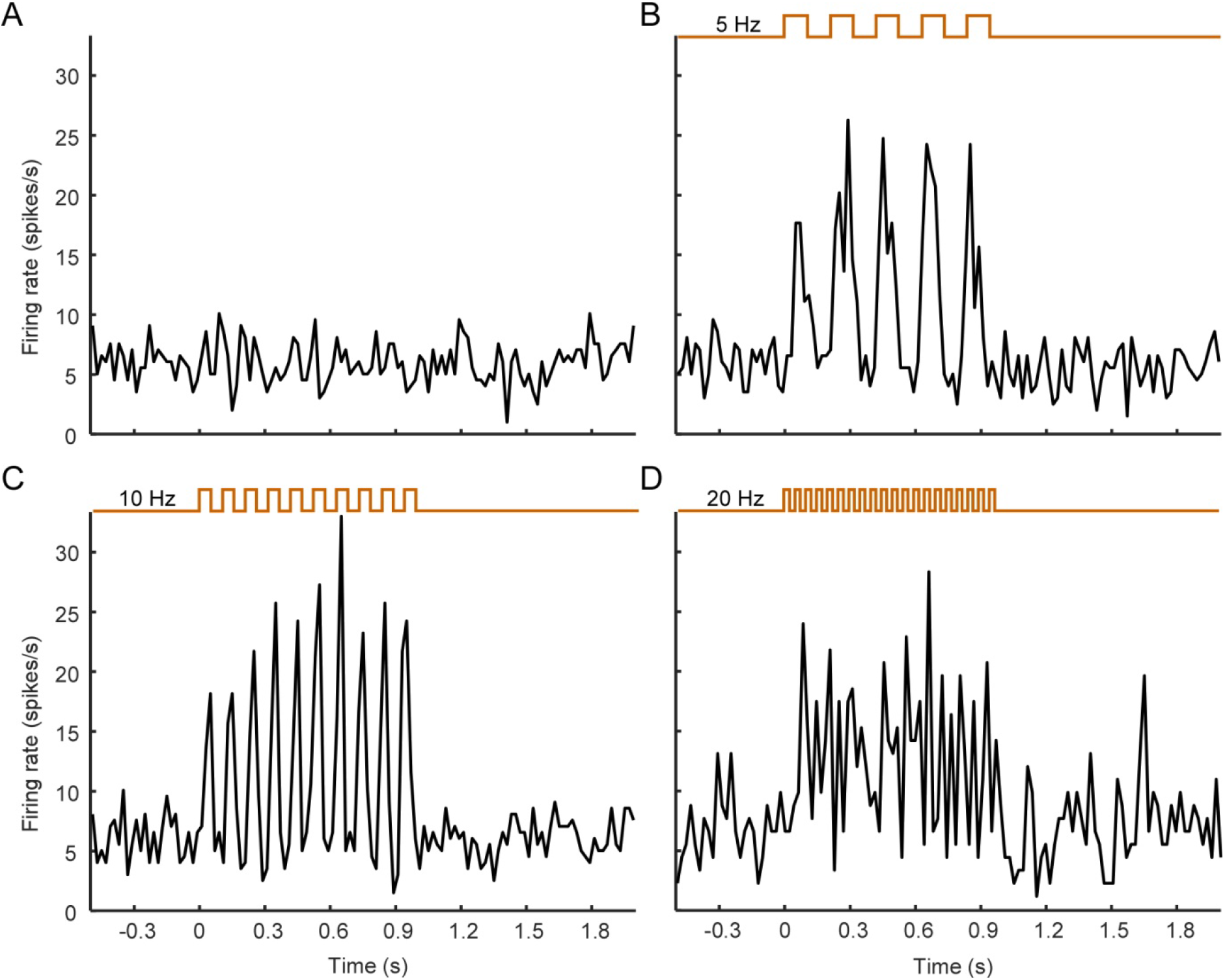
Excitation caused by 594 nm wavelength laser in cortical cells. Peri-stimulus time histogram with no laser activation (a), 5 Hz (b), 10 Hz (c), and 20 Hz (d) square wave laser stimulation. The cell response follows the laser pulses at 5 and 10 Hz, but breaks down at 20 Hz although the firing rate is still elevated above baseline. 20 mW at fiber tip.

To test the effectiveness of our virus in manipulating long-range connections, and to explore the causal effects of pulvinar activation and inhibition on cortical activity, we tested cortical responses to visual stimuli during laser manipulation of LP following V2 injections (Figures 6 and 7). V2 was chosen because, as we have recently shown, pulvinar sends many more projections to higher visual cortex than to V1 (Scholl et al., 2020). Local field potential (LFP) and spiking activity were recorded during presentation of flash stimuli over six cortical penetrations and LP optic fiber sites. Visually evoked potentials (VEP) and mean firing rates (MFR) were measured in response to the onset of a white screen following periods of black (Figures 10 and 11). Offset responses were also recorded but not considered for analysis since typical VEPs and MFR were much larger at the stimulus onset (data not shown).

**Figure 6.**
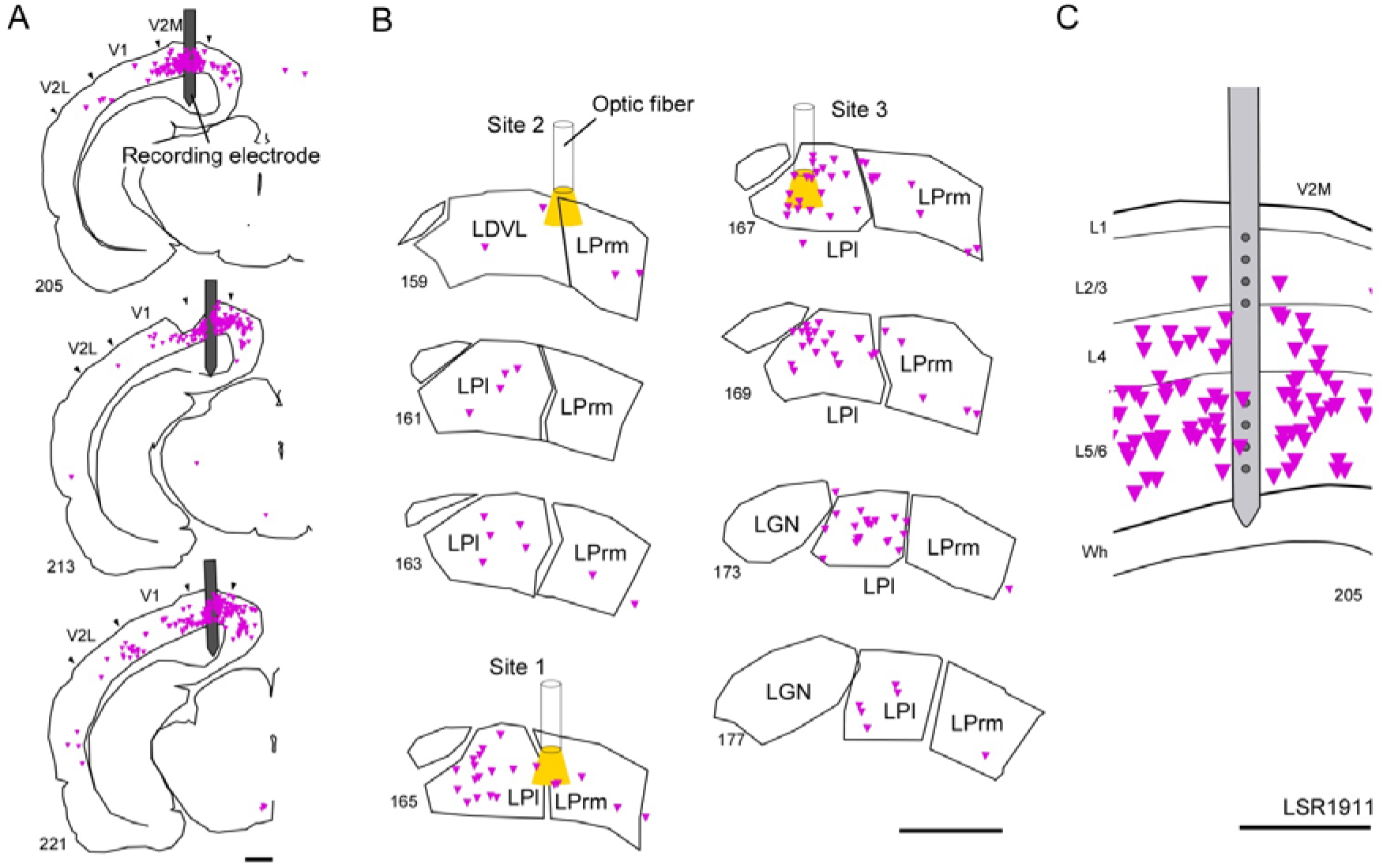
Recording from medial V2 neurons while optically manipulating pulvinar input. Reconstruction of case LSR1911. mScarlet-STGtACR2-STChrimsonSA rabies virus was injected into medial V2 (V2M; panel a) and retrogradely infected cells in LPl and LPrm (b). Recordings were made across several cortical layers at the injection site (c) while simultaneously activating or inhibiting cells in LP with 594 nm and 473 nm wavelength laser light in order to uncover thalamocortical modulation of activity in V2M. Three sites were targeted sequentially in LP. Site 1 targeted central LP, Site 2 targeted anterior LP, and Site 3 targeted LPl only. Scale bars in (a) and (b) equal 1 mm; scale bar in (c) equals 0.5 mm.

**Figure 7.**
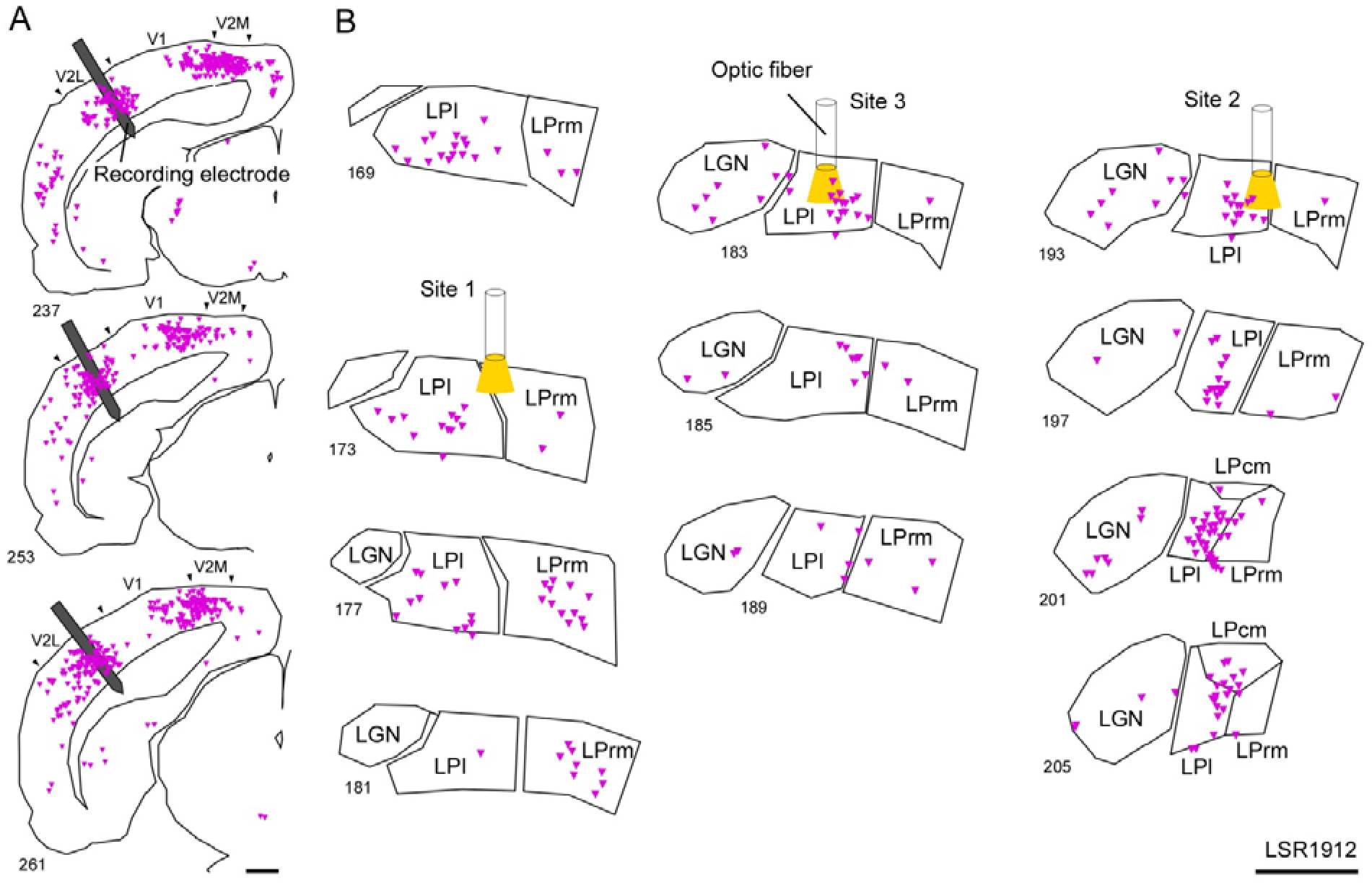
Recording from lateral V2 neurons while optically manipulating pulvinar input. Reconstruction of case LSR1912. Virus was injected into lateral V2 (V2L, panel a), retrogradely infecting neurons in LPl, LPrm, and LPcm (b). An optic fiber was lowered sequentially into three sites targeting LPl during extracellular recordings of cells in V2L. Scale bars equal 1 mm.

At several V2 recording sites, changes in VEP amplitude were immediately noticeable following both activation (594 nm laser) and inactivation (473 nm laser) of pulvinar neurons (Figure 8a, 8c). In these channels, activation of pulvinar lowered evoked amplitude, whereas inactivation raised amplitude. Other recording sites showed signs of amplitude change in one direction but not the other as shown in Figure 8b, in which inactivation (blue laser) caused increased amplitude whereas activation caused no change. Importantly, no channels showed amplitude increase following pulvinar activation or amplitude decrease following pulvinar inactivation.

**Figure 8.**
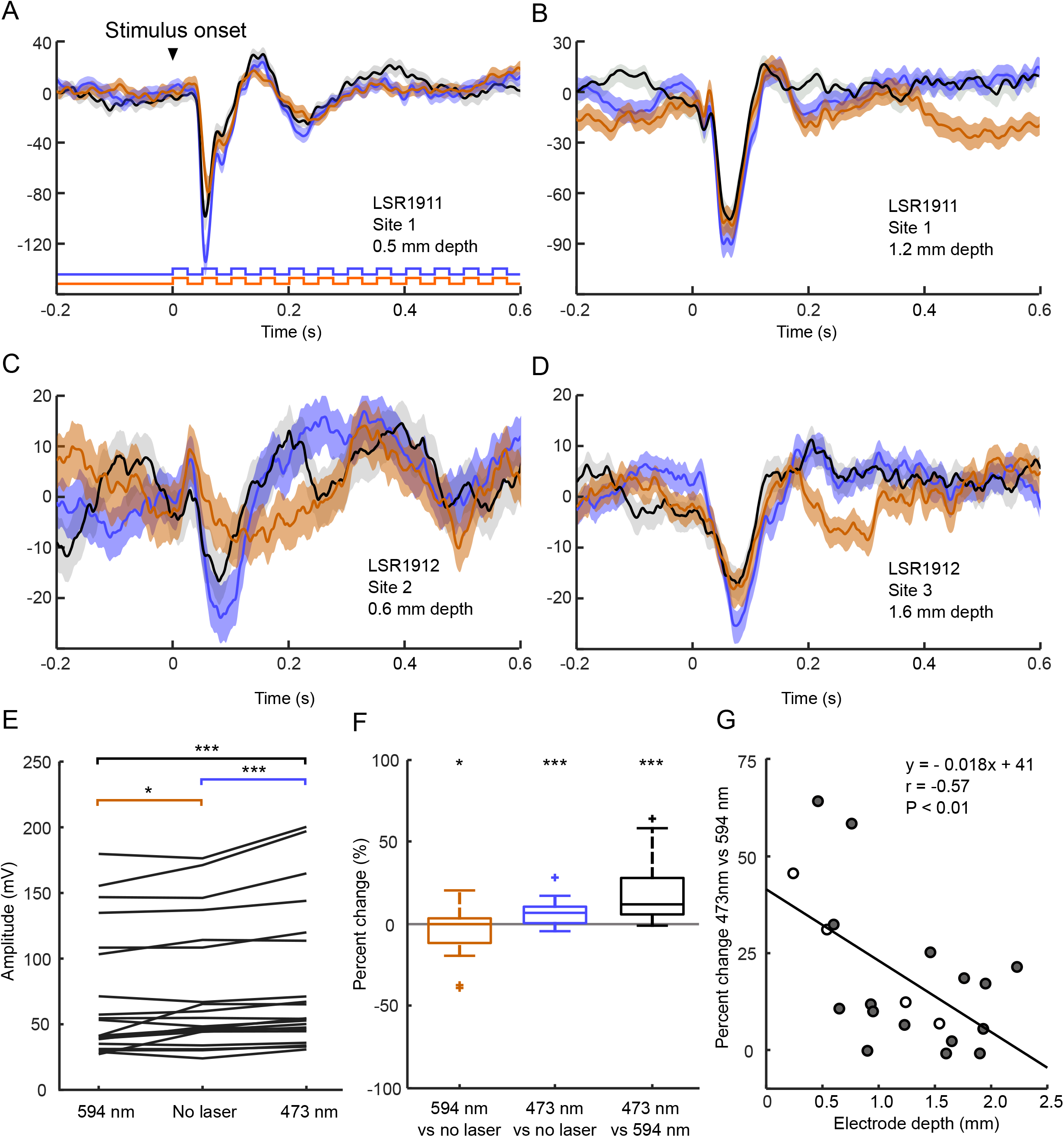
Visually evoked potentials (VEP) in V2 during pulvinar manipulation of mScarlet-STGtACR2-STChrimsonSA rabies virus infected cells. (a-d) Examples of VEP during no laser (dotted lines), yellow laser (orange lines) or blue laser (blue lines) stimulation of LP. (e) Population amplitude of VEPs in V2 when LP neurons were activated by yellow (594 nm) laser, when no laser was used, and when LP neurons were suppressed by blue (473 nm) laser. Evoked responses in V2 are strongest when LP neurons are inactivated and weakest when LP neurons are activated. (f) Percent change between activation, suppression, and no laser conditions. (g) Comparison of electrode depth and percent change between suppression and activation of LP neurons. Sites recorded in V2L are shaded; sites recorded in V2M are open circles. Solid line indicates a significant linear correlation between electrode depth and strength LP manipulation had on cells in V2 at that depth.

On average, amplitude in response to stimulus onset was significantly lower during periods of LP activation (594 nm laser) compared to baseline (Figure 8e-f; n = 20, *P* < 0.05), and significantly higher for periods of LP inactivation (473 nm wavelength laser) compared to baseline (*P* < 0.001, Wilcoxon sign test). Comparing the two laser wavelengths directly, there was a significantly higher amplitude for 473 nm wavelength light than for 594 nm wavelength light (*P* < 0.001). Without any visual stimulus, neither laser wavelength caused significant changes to amplitude.

The difference in amplitude between the two laser wavelengths essentially measured the strength LP manipulation had on a particular cortical recording site. We compared this manipulation strength against electrode depth to identify which cortical layers were most affected by pulvinar. Recording sites in superficial cortical layers were the most strongly manipulated. Across the whole population, electrode depth was negatively correlated with change in amplitude between the two laser wavelengths (r = −0.57, *P* < 0.01; see Figure 8g).

Cortical single unit responses to the same stimuli were also affected by pulvinar manipulation. Three V2 neuron examples are shown in Figure 9a-c. Firing rates were significantly lower on average during pulvinar activation compared to without modulation (Figure 9d-e; n = 28, *P* < 0.05), and significantly higher during pulvinar inactivation (473 nm laser) versus pulvinar activation (594 nm; *P* < 0.05). Cortical electrode depth was also significantly correlated with firing rate change (r = −0.35, *P* = 0.05).

**Figure 9.**
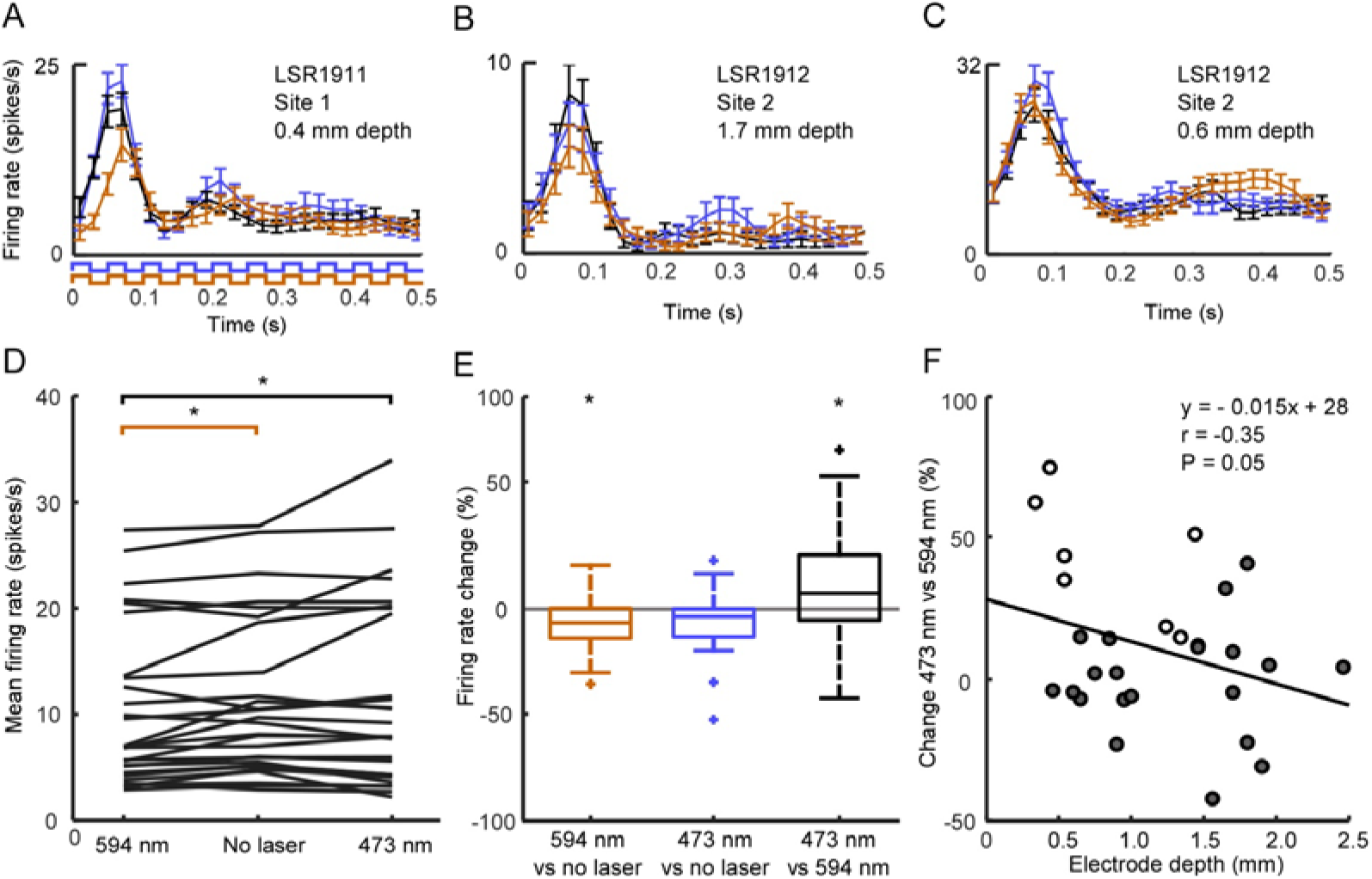
Single unit responses to flash stimulus with and without laser manipulation of pulvinar. (a-c) Three example cells with varying degrees of modulation from pulvinar. Colors indicate which laser manipulation was used, while dotted lines indicate no manipulation was used. (d) Population response during pulvinar stimulation with yellow laser (594 nm), no laser, or blue laser (473 nm) light. (e) Percent change over no laser condition for yellow and blue lasers, and percent change from yellow laser to blue laser conditions. (f) Comparison of electrode depth to percent change in firing rate between blue laser and yellow laser. Open circles indicate units were recorded from V2M while shaded circles indicate units were recorded from V2L.

Cells in V2 were also tested for responses to sinusoidal drifting grating stimuli at different spatial frequencies, temporal frequencies, orientations, and sizes. A subset of these cells (n = 8) were tested for size preference during laser stimulation of pulvinar. These cells were highly selective to the diameter of the stimulus, with optimal responses at sizes ranging from 18° to 40° (*M* = 26 ± 7°) and maximally suppressed at a size ranging from 50° to 80° (*M* = 63 ± 12°).

As shown by the example cells (Figure 10a and b), mean V2 cell firing rates at the optimal diameter were significantly lower when pulvinar was inactivated (473 nm laser) than without pulvinar modulation (*P* < 0.05; Figure 10c). No significant change in firing rate at the optimal size was observed during pulvinar activation (594 nm laser). In contrast, at the maximally suppressed size, cells in V2 were more clearly affected by pulvinar activation than inactivation (see example in Figure 10a). On average, firing rates at the suppressed size increased significantly during pulvinar activation (594 nm laser; *P* < 0.05). No significant change in firing rate at the suppressed size occurred during pulvinar inactivation (473 nm laser).

**Figure 10.**
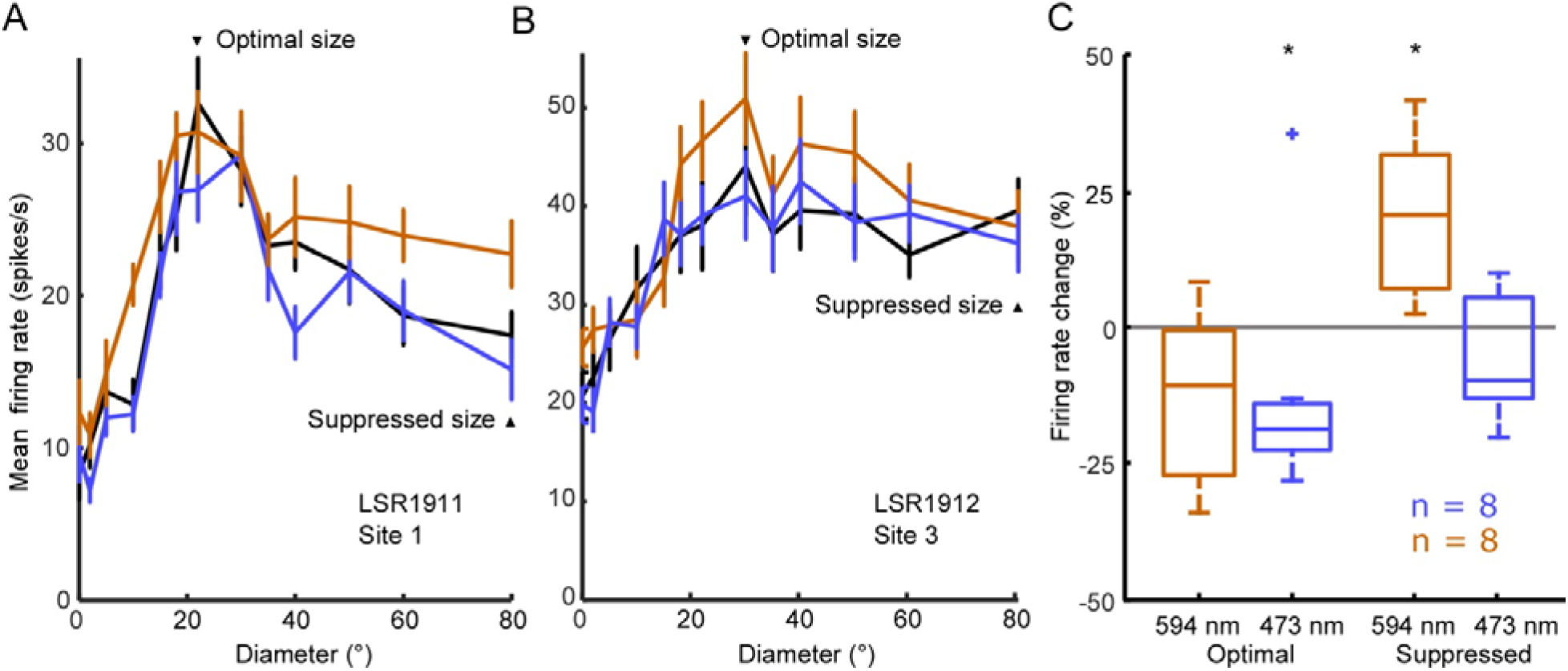
Changes in cortical size tuning due to optical manipulation of pulvinar. Two examples of size tuned cells in rat V2, one with decreased firing rate at optimal size (20°) due to pulvinar inactivation and increase firing rate at the maximally suppressed size (80°) due to pulvinar inactivation (a), and one with increased firing rate at optimal size due to pulvinar excitation (b). Population medians and quartiles of percent change at the optimal diameter (c) show that inactivation of pulvinar consistently decreased firing rate.

## Discussion

Here we demonstrated bi-directional optical control over populations of cortical and thalamic neurons using a modified rabies virus. To our knowledge it is the first example to achieve independent excitatory and inhibitory control of a neuron *in vivo* and with this virus. Using this new tool we showed that pulvinar neurons suppress activity in higher visual cortex in the rat during flash stimulus presentation, and that pulvinar has influence over cortical size tuning.

Overall, our method of dual opsin delivery is extremely efficient, utilizing the best currently available light-sensitive ion channels, fluorescent protein, and optimized rabies virus vector. Previous attempts to deliver bimodal opsins *in vivo* using separate lentiviral vectors (Zhang et al., 2007), or transgenic animals (Yizhar et al., 2011) have not been able to target the same population of cells with both transgenes. Our method using a rabies virus vector, however, is capable not only of simultaneous expression of both opsins in all infected cells, but also of selectively infecting populations of projection neurons to the injection site without the need for transgenic animals (Ghanem & Conzelmann, 2016). An alternative approach is to perform separate experiments using two or more optogenetic approaches, but again this is limited to separate populations of neurons, not only slowing research due to the increased requirement for animals and surgical procedures, but also limiting the sophistication of excitation/inhibition modulations of the neuron population.

Several improvements to the virus could be beneficial. First, the relationship between laser power and change in firing rate has not fully been quantified. We simply used the maximum available yellow laser power and 25% maximum available blue laser power after our initial tests showed that this was a suitable combination in most cases. However, we are uncertain whether interactions between the opsins could be diminishing the potential efficiency of either protein. The power range within which to induce only excitation or inhibition in our design is large, due not only to spectral separation, but also relative sensitivity and differential expression levels of ChrimsonSA and GtACR2. Additional testing, preferably *in vitro* using patch clamped cells and direct laser activation and inactivation, could be used in future studies to more precisely determine the optimal power range. Second, although useful for filling whole cells and visualizing cells *in vivo* due to the bright mScarlet, the fluorescence in axons is so bright that finding cell bodies can be challenging (see Figure 2). This could be overcome by fusing the fluorescent protein to one or both of the channelrhodopsins, limiting fluorescence to the cell soma at the membrane.

Pulvinar inactivation has been studied in the past by methods other than what we described here. For example, superior colliculus lesions in mice cause changes to speed tuning across multiple cortical areas (Tohmi et al., 2014). However these changes are hard to attribute to the pulvinar since superior colliculus projects to the lateral geniculate nucleus and other thalamic nuclei (Partlow, et al., 1977; Pasquier & Villar, 1982). Pulvinar inactivation by GABA receptor agonist injections drastically lowered cortical firing rates in monkeys (Purushothaman et al., 2012; Zhou et al., 2016), but such injections cause major changes to thalamic physiology, as well as potentially causing damage to cortex (Lomber, 1999; Majchrzak & Di Scala, 2000). Other nondestructive methods have also been used to correlate task-dependent activity in the pulvinar to activity in cortical areas (Saalmann et al., 2012), but these lack causal inference. Our method of optically manipulating smaller, targeted networks of cells shows that pulvinar can both increase and decrease firing rates in response to visual stimuli.

Pulvinar activation and inactivation had different effects on different systems, further illustrating the need for bi-directional control of membrane potentials. During flash stimulus presentation, activation of pulvinocortical projections caused decreased evoked potential amplitudes and firing rates, yet size tuned cells were facilitated at large diameters by pulvinar activation. Pulvinar projections to cortex do synapse with multiple cortical layers in rats (Nakamura et al., 2015), it’s possible these different contextual effects are mediated by different laminar networks in the cortex, one inhibitory and one excitatory. We did observe laminar specificity of flash modulation, as the effect of pulvinar manipulation was greatest in superficial layer cortical cells (see Figures 8g and 9f), suggesting that the pulvinar modulates feedforward projections (Felleman & Van Essen, 1991; D’Souza & Burkhalter, 2017). At least for simple flash stimuli, pulvinar neurons may amplify responses traveling through hierarchical cortical areas to stimuli inside their receptive fields. Future studies could determine whether layer-specific modulation occurs for more complex stimuli, or during behavioral tasks in awake animals. For example, neurons in the dorsomedial primate pulvinar that are enhanced during covert attention (Petersen et al., 1985) might modulate responses in cortex without the need for optogenetic manipulation.

The experiments here targeted mainly the lateral portion of the pulvinar, because in our initial injections most of the projections to V2 originated from the lateral subdivision, LPl (see Figure 2d). However, all of the three rodent pulvinar subdivisions send projections to visual cortex, although each has a distinct pattern of connectivity (Nakamura et al., 2015; Scholl et al., 2020). Differences between the subdivisions could lead to functional differences as well. For example, the caudal portion of the pulvinar receives superior colliculus input while the rostromedial portion does not (Takahashi, 1985), and activity in the caudal pulvinar in mice is dependent on superior colliculus while activity in rostral pulvinar is dependent on cortex (Bennett et al., 2019). To test whether or not neurons in each subdivision have similar effects on visual cortex, future experiments could simply place the optic fiber carefully in each subdivision otherwise using the same experimental paradigms shown here.

Cortical areas may also receive differential projections from the pulvinar. We previously made injections into both medial and lateral portions of V2, expecting differences in projection strength or cortical modulation by pulvinar neurons (Scholl et al., 2020). However, roughly the same number of neurons were found to project to both V2 areas, consistent with previous work (Nakamura et al., 2015), and the effects of optical control over pulvinar neurons were consistent across V2M and V2L (see Figures 8g and 9f). Specializations within rodent V2 (Tohmi et al., 2014; Nishio et al., 2018) and of projections between pulvinar and V2 (Juavinett et al., 2019) suggest that the pulvinar supports multiple channels of transthalamic computation; further research in this area is warranted.

The virus presented here represents a powerful new tool for studying the causal relationship of neural populations within and between neural structures. Models of neural systems can be tested more rigorously by taking advantage of the combined benefits of retrograde expression, bright fluorescences, bidirectional optogenetics, and fast kinetics of ChrimsonSA allowing for more sophisticated manipulation of long-range neural circuitry.

